# Subjective visibility report is facilitated by conscious predictions only

**DOI:** 10.1101/2020.07.08.193037

**Authors:** Josipa Alilović, Heleen A. Slagter, Simon van Gaal

## Abstract

Predictions in the visual domain have been shown to modulate conscious access. Yet, little is known about how predictions may do so and to what extent they need to be consciously implemented to be effective. To address this, we administered an attentional blink (AB) task in which target 1 (T1) identity predicted target 2 (T2) identity, while participants rated their perceptual awareness of validly versus invalidly predicted T2s (Experiment 1 & 2) or reported T2 identity (Experiment 3). Critically, we tested the effects of conscious and non-conscious predictions, after seen and unseen T1s, on T2 visibility. We found that valid predictions increased subjective visibility reports and discrimination of T2s, but only when predictions were generated by a consciously accessed T1, irrespective of the timing at which the effects were measured (short vs. longs lags). These results further our understanding of the intricate relationship between predictive processing and consciousness.

## 1. Introduction

Within the predictive processing framework (Clark, 2013; de Lange et al., 2018; Friston et al., 2009; Hohwy, 2013; Rao, 2005; Rao & Ballard, 1999) perception is understood as resulting from an interaction between bottom-up stimulus-related neural activation and top-down prediction-driven biases. In this framework, the brain continually formulates hypotheses (predictions) based on past experience about the hidden causes of its sensory input (Friston, 2009; Seth, 2014, Hohwy, 2013). Such sensory predictions constrain possible interpretations and shape perception (de Lange et al., 2018; Hohwy, 2013; Panichello et al., 2013). It has for example, been shown that predictions facilitate object recognition (Bar, 2003; Bar et al., 2006; Chaumon et al., 2014; Oliva & Torralba, 2007; Chang et al., 2016; Melloni et al., 2011; Stein & Peelen, 2015) and scene perception. Predictions may especially benefit perception when the perceptual context is ambiguous, involves competing alternatives (Denison, Piazza, & Silver, 2011; Pinto, Gaal, Lange, Lamme, & Seth, 2015) or when sensory input is weak (Summerfield et al., 2006).

Predictions have also been shown to increase the likelihood of conscious report of visual stimuli (Meijs et al., 2019; Meijs et al., 2018). This was illustrated in an attentional blink (AB) paradigm in which two targets (T1 and T2) are presented within a rapid stream of distractors. In this task, conscious access of T2 after correct T1 identification is significantly decreased when T2 follows T1 after approximately 200-500 ms compared to when T2 follows T1 after a longer temporal interval (Raymond et al., 1992), a phenomenon called the ‘attentional blink’. Interestingly, Meijs et al. (2018, 2019) showed that when the identity of T1 predicted T2 identity, valid predictions decreased the AB to T2 (or in other words valid predictions increased the likelihood of T2 identification). This effect was furthermore shown to depend on T1 visibility (Meijs et al., 2018). That is, T1 needed to be consciously accessed in order for its predictive nature to affect subsequent conscious access of T2. These findings suggest that 1) predictions impact conscious access, and 2) that these predictions need to be implemented consciously to be effective. Building on these findings, the current study aimed to address two open issues concerning the nature of the relationship between predictions and conscious access. First, we examined how predictions modulate conscious access, using objective discrimination performance measures and subjective visibility reports. Second, we further explored the temporal profile of prediction effects, and whether non-conscious predictions may be able to modulate conscious report when the weaker and more fleeting nature of non-conscious processing is taken into account.

Regarding the first question, a significant body of work suggests that prediction effects are perceptual in nature. Studies have reported that predictions modulate sensory responses (e.g. Kok, Rahnev, et al., 2012; Richter et al., 2018), improve sensitivity of early feature-detectors (Teufel et al., 2018) and sharpen neural representations in sensory brain regions (Kok, Jehee et al., 2012; Yon et al., 2018). Moreover, a recent study by Gandolfo & Downing (2019) provided evidence for a causal link between prediction-related neural activity and perceptual outcomes. The authors applied fMRI-guided online transcranial magnetic stimulation (TMS) over two category-selective brain regions, the extrastriate body area (EBA) and the occipital place area (OPA), during the presentation of a prediction cue that validly or invalidly predicted a body or a scene feature on images participants needed to judge. Strikingly, TMS over the task-relevant site (EBA during the body task, OPA during the scene task) eliminated the validity effect by the prediction cue. However, others have suggested that predictions (in some cases) may simply change the decision criterion (Bang & Rahnev, 2017) and modulate metacognitive judgements related to perceptual decisions (Sherman et al., 2015). For instance, Bang and Rahnev (2017) contrasted the effects of probabilistic pre- and post-stimulus cues on stimulus sensitivity and decision criterion (d’ and c from the signal detection theory). They reasoned that pre-stimulus cues can affect both the sensory signal and the decision criterion, while post-stimulus cues, by virtue of being presented after the sensory signal, can only affect the decision criterion in case when the stimulus is masked (the authors used backward masking to interrupt sensory reverberations of stimuli to eliminate the possibility that prediction cues could retroactively enhance perception, Sergent et al., 2013). The authors found that post-cues influenced participants’ decision criterion (c) to a larger extent, while there was no difference between pre and post cues in their effect on stimulus sensitivity (d’). Furthermore, the effects of pre and post cues on the decision criterion were found to be highly correlated, suggesting that both cues act through similar mechanisms. This may suggest that predictions modulate the decision criterion, such that, for instance, a stimulus category is reported more readily when it is congruent with the probabilistic structure of the task. In line with this, other studies have reported that valid predictions did not modulate early sensory responses to stimuli while they did increase the likelihood of correct identification (Alilović, et al., 2019; Meijs et al., 2019; Rungratsameetaweemana et al., 2018). Meijs et al. (2018) also aimed to arbitrate between prediction-induced decision criterion shifts versus perceptual shifts. In that study (specifically experiment 1), the authors showed that valid predictions even increased the likelihood of conscious report when subjects only had to report whether they had seen a T2 or not (seen/unseen response), irrespective of T2 identity. As this dependent measure renders the identity of T2 irrelevant, there is no obvious decision bias that may affect this response, suggesting that predictions may be able to induce perceptual shifts (Meijs et al., 2018). Yet, it could still be that when prediction and target identity matched this lowered the decision threshold for reporting “seen”, while the true experience of the target remained unaffected. Further investigating whether predictions modulate the perceptual experience of stimuli and/or whether they may primarily change the criterion to report it, was the first goal of this study. To that end, in Experiments 1 and 2, we measured prediction effects on perceptual experience using the Perceptual Awareness Scale (Overgaard et al., 2006; Sandberg et al., 2010), which allowed us to express the effects on a more fine-grained scale that focuses on subjective visibility.

Regarding our second question, Meijs et al. (2018) suggested a tight link between consciousness and predictions, because only consciously perceived T1s (the prediction cues) were observed to facilitate subsequent conscious access of T2. Yet, it is a topic of an ongoing debate whether, or in which circumstances, predictive processing is automatic versus “strategic” in nature (Grotheer & Kovács, 2016). Some predictions, such as those pertaining to contextual and environmental associations on natural scenes, which we acquire through life-long learning, seem to be more hardwired and therefore less dependent on consciousness (e.g. Brodski et al., 2015; see also O’Callaghan et al., 2017; Chang et al., 2016; Grotheer & Kovács, 2016). Moreover, low-level violations of predictions are registered by the brain non-consciously (Kimura & Takeda, 2015; King et al., 2013; Meijs, et al. 2018), as for example illustrated by the presence of mismatch responses in auditory oddball tasks in patients with disorders of consciousness, such as the vegetative state (Bekinschtein et al., 2009; for a review see Morlet & Fischer, 2014). Lastly, it is generally known that non-conscious effects are much weaker and shorter-lived than their conscious counterpart (Dehaene et al., 2006; Greenwald et al., 1996; van Gaal & Lamme, 2012; Van Vugt et al., 2018). Together, these observations raise the possibility that non-conscious predictions can affect perception, but that their effects may be only detectable on a sufficiently short temporal scale. For instance, a task design that includes short temporal lags between two targets that are predictive of one another, could be more sensitive in observing faint and quickly decaying non-conscious prediction effects (cf. Meijs et al., 2018). In Experiments 2 and 3, we tested this possibility using an attentional blink task in which T2 could follow T1 directly or after only one distractor (i.e., at lags 1 and 2). In the Meijs et al. study, T2 followed T1 relatively late, after two distractors (i.e., at lag 3).

In summary, in three behavioral experiments, we address two outstanding questions concerning the relationship between consciousness and predictions, namely the perceptual or post-perceptual (or both) nature of prediction effects on perceptual experience, and whether non-conscious predictions may modulate perceptual conscious report when the quickly-decaying nature of non-conscious processes is taken into account.

## 2. Experiment 1

### 2.1 Materials and Method

In Experiment 1, we directly tested whether predictions may affect subjective visibility reports of stimuli. To that end, we employed an attentional blink task in which, on each trial, T1 identity predicted which T2 letter was more likely (cf. Meijs et al., 2018). Participants rated their subjective perceptual awareness of the target using the PAS.

#### 2.1.1 Participants

Thirty-two students, recruited from the University of Amsterdam, participated in Experiment 1. This sample size was chosen based on a study by Meijs et al. (2018) (Experiment 1), which showed a significant effect of conscious prediction on T2 identification (η^2^=0.23) using a similar sample size of n=26. All participants were right-handed, reported normal or corrected-to-normal vision, and no history of a psychiatric or neurological disorder. All participants gave written informed consent prior to the start of the experiment and received research credits or money (10 euros per hour) for compensation. One participant was excluded from the analysis due to missing data on the PAS scale. The final sample thus consisted of 31 participants (19 female, mean age=20.34 years, SD=1.76). The study was approved by the ethical committee of the Department of Psychology of the University of Amsterdam.

#### 2.1.2 Materials

All stimuli were generated using Matlab and Psychtoolbox-3 software (Kleiner et al. 2007) within a Matlab environment (Mathworks, RRID:SCR_001622). Stimuli were presented on 1920×1080 pixels ASUS VG236H monitor at a 120-Hz refresh rate on a “black” (RGB: [0 0 0], ± 3 cd/m^2^) background and were viewed with a distance of ∼ 70 cm from the monitor.

#### 2.1.3 Task and Design

The task was adapted from Meijs et al. (2018). In total, 17 uppercase letters, excluding I, L, O, Q, U and V, were shown in a rapid serial visual presentation (RSVP) stream on each trial of the AB task (Fig. 1A; cf. Meijs et al., 2018). Letters were presented at fixation in a monospaced font (font size: 40 points, height ∼1.2°), in white RGB: [255 255 255]; 320 cd/m^2^) for 92 ms per letter. A letter was never repeated on a given trial. The first target letter (T1: G or H, presented with equal likelihood) was presented on 90% of all trials, always as the fifth letter in the RSVP. T1 was presented in green RGB: [0 255 0]; ± 320 cd/m^2^), and surrounded by four small gray squares (RGB: [200 200 200], ± 188 cd/m^2^; size: 0.35°; midpoint of each square centered at 1.30° from fixation) (Fig 1A). The second target letter (T2: D or K) followed T1 on 80% of trials. Like T1, T2 was always surrounded with placeholders (four small gray squares). On 20% a distractor letter (any letter other than T1/T2) was presented instead of a T2, also surrounded by gray squares. Thus, on trials without T1 or/and T2, a random distractor letter was presented surrounded with placeholders instead. As in Meijs et al., T2s were presented at lags 3 and 10, which corresponded to 275 and 917 ms of temporal separation between targets, respectively. Each lag was equally likely.

**Figure 1.**
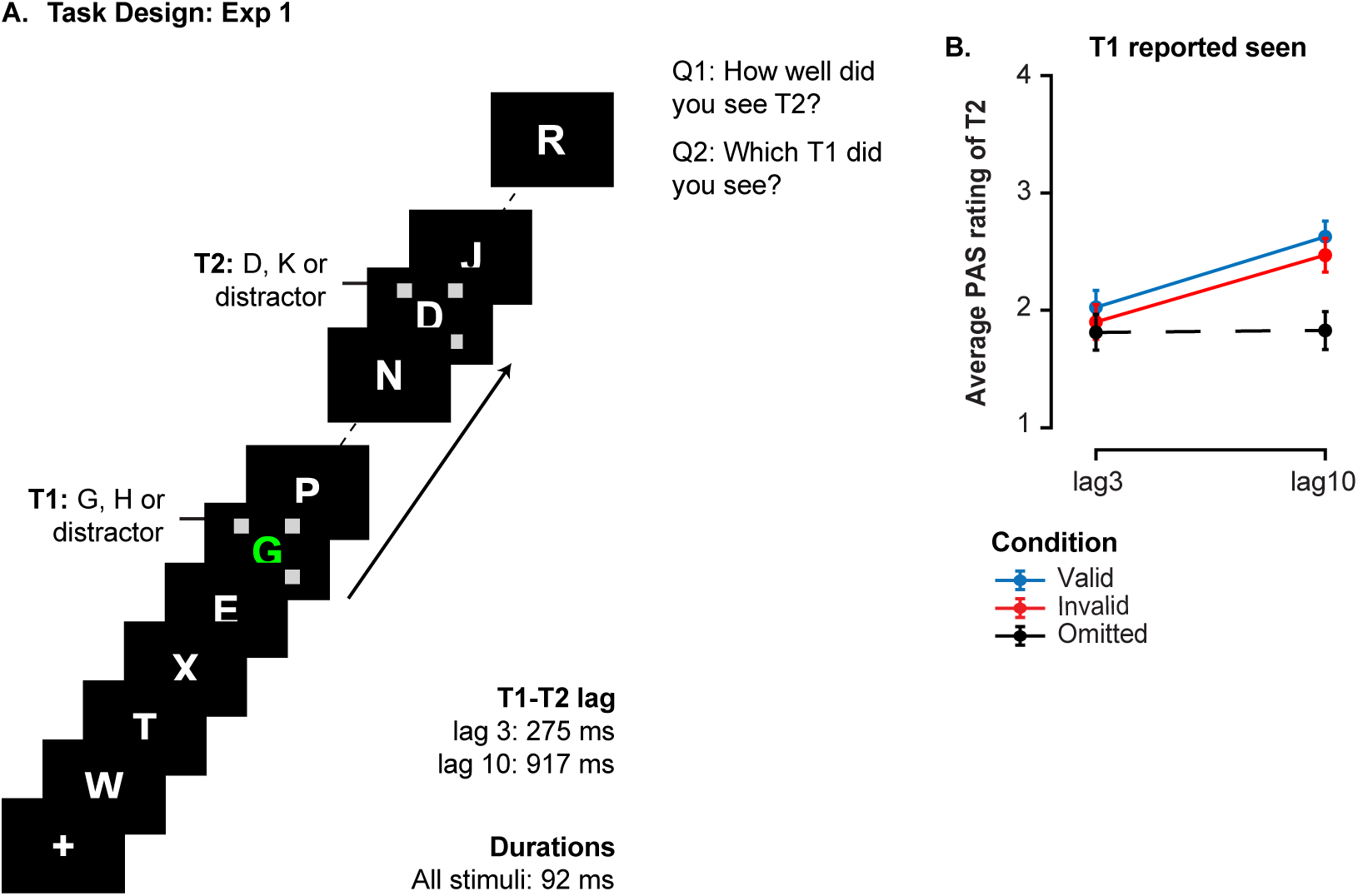
Task design and behavioral results of Experiment 1. **A)** The trial structure of the AB task. On each trial, we showed an RSVP stream of letters in which two predefined target letters were embedded. The first target (T1: a green G or H) always appeared at the fifth position in the stream surrounded by placeholders. The second target letter (T2: D or K) was shown at one of 2 lags (lags: 3 and 10) after T1, also surrounded by placeholders. On 20% of the trials, a random distractor letter was presented instead of T2 (T2 omitted trials). The identity of T1 predicted the identity of the T2 target letter with 75% validity, which introduced validly and invalidly predicted T2s. At the end of each stream, participants provided two answers. First, they rated the perceived visibility of T2 using a 4-point scale (PAS; 1-No experience; 2-A vague experience; 3-An almost clear experience; 4-A clear experience). Thereafter they reported the T1 letter they had seen. **B)** Average PAS rating of T2s at each T2 lag for validly and invalidly predicted T2s and for T2 omitted trials on which T1 was correctly identified (T1 seen). Error bars represent SEM. Validly predicted T2s were subjectively experienced as more visible, independent of lag.

Importantly, the identity of T1 letter predicted the identity of T2 letter and participants were informed about this relationship, as in Meijs et al. For instance, after seeing G as the T1 letter, participants were informed that, if a T2 was presented, the letter D would be presented in 75% of cases, while the letter K would be less likely (25%). The mapping of T1s and T2s was counterbalanced across participants such that G-D and H-K combinations were more likely for odd participant numbers, and G-K and H-D for even participant numbers. Each RSVP stream ended with a 150 ms blank period, after which participants gave their responses. Critically, in order to minimize the contribution of a response bias towards reporting the more frequent T2 letter after a given T1 letter (e.g., D follows G) and to measure the subjective visibility of T2 instead, participants were asked to give a rating of their perceptual experience of T2 on a 4-point Perceptual Awareness Scale (PAS; 1-No experience; 2-A vague experience; 3-An almost clear experience; 4-A clear experience) (Overgaard et al., 2006; Sandberg et al., 2010). Following the T2 rating, participants were also asked to identify which T1 letter they had seen. They were instructed to give rating 1 when they did not see T2 and to refrain from entering a letter when they did not see T1. The responses were confirmed by pressing the spacebar on the keyboard or when a timeout of 4 seconds had passed. The inter-trial interval was 300-500 ms.

#### 2.1.4 Procedure

The experiment consisted of one 2-hours-long session. After providing consent, participants first read the instructions and completed 3 practice blocks, each consisting of 40 trials. The experimenter also verbally instructed participants, reiterating the most important aspects of the task, in particular the predictive mapping between T1 and T2. The main task consisted of 15 experimental blocks, each consisting of 40 trials. Each block was followed by a self-paced break, during which participants received feedback about their performance (percentages of T1 misses and T1 false alarms). After every four blocks, there was a longer forced break during which the experimenter went into the testing cubicle to assure that the participant took a break. The experimenter would also check on participants’ performance in identifying T1 (percentages of misses and false alarms) and encourage them to keep their performance up to high levels.

### 2.2 Data Analysis

The main statistical analysis tested whether the average T2 visibility rating, expressed on the PAS scale, differed significantly between the two prediction conditions (valid and invalid) and whether this difference was modulated by the lag at which T2 was presented with respect to T1 (3 and 10). Trials on which no T1 response was given when T1 was actually presented (3.98% of all trials) or an impossible T1 response was given (two T1 letters reported on 0.08% of all trials), or trials on which no or an impossible T2 rating was given (two or more T2 ratings or a rating > 4; in total, 5.47% of trials) were excluded from the analysis. To compute average T2 PAS ratings for the two prediction conditions, separately at lag 3 and 10, we used only T1-correct trials, i.e. T1-present trials in which T1 was correctly identified. Using T1-correct trials, we also computed the average T2 PAS rating for trials in which T2 was omitted, separately at lag 3 and 10.

Average PAS ratings were entered into a 2 × 2 repeated measures ANOVA with the within-subject factors Prediction (valid, invalid) and Lag (lag 3, lag 10) to evaluate whether valid versus invalid predictions modulated subjective visibility reports of T2 as a function of lag. Significant main and interaction effects were followed-up by paired-sample *t*-tests. All effects were followed up by a Bayesian equivalent of the same test, which quantified the strength of evidence for the null hypothesis (H_0_) (Wagenmakers et al., 2018). Bayes factors (BF_01_) were computed using JASP’s (JASP Team, 2018) default Cauchy prior. We conducted the Bayesian equivalent of the repeated measures ANOVA with the same within-subject factors as in the classical repeated measures ANOVA and the Bayesian equivalent of the paired-sample *t*-tests to determine the strength of evidence in favor of the H_0_ for the main and interaction effects of interest, and simple comparisons, respectively. In case we needed to quantify evidence for an interaction effect, we computed exclusion Bayes factor (BF_excl_) across matched models, which indicates the extent to which data supports the exclusion of an interaction effect, taking all relevant models into account. In accordance with terminology suggested by Jeffreys (1961), we labeled the effect sizes from 1 to 3 as anecdotal evidence in favor of H_0,_ values from 3 to 10 as substantial, and those above 10 as strong evidence in favor of H_0_.

### 2.3 Results

On average, participants achieved high performance in identifying T1 (mean=88.6%. SD= 8.77%, T1 false alarm rate was 3.33%). Average T2 PAS ratings in each prediction condition and each lag were statistically compared in repeated measures ANOVAs. This analysis revealed that predictions as expected modulated T2 PAS ratings (main effect of prediction: F_1,30_=4.58, p= .04, η^2^ =0.132, BF_01_=1.94, Figure 1B). Validly predicted T2s were rated higher than invalidly predicted T2s. Furthermore, we found a clear AB pattern (participants rated T2s presented at lag 3 lower on the PAS scale than T2s at lag 10, F_1,30_=31.84, p<.001, η^2^=0.515, BF_01_=1.05×10^−9^), but no interaction between Lag and Prediction (F_1,30_=0.24, p=.63, η^2^=0.008, BF_excl_=3.59). These findings indicate enhanced subjective visibility reports for predicted (compared to unpredicted) targets both in and outside the AB time window.

The results of this experiment highlight that conscious predictions can modulate subjective visibility reports of events. Validly predicted T2s were reported to be experienced more clearly than invalidly predicted ones, both within and outside the AB time window. These results corroborate earlier findings (e.g. Meijs et al., 2018; 2019) using a report measure that is likely less sensitive to response biases and further strengthen the conclusion that predictions may lead to shifts in the conscious reportability of visual stimuli.

## 3. Experiment 2

Results of our Experiment 1 showed that changes in subjective visibility reports as a consequence of learned predictions can be measured on the PAS scale. The PAS scale is arguably less likely influenced by response biases, i.e. tendency towards reporting (or guessing) the more probable stimulus category, as opposed to a discrimination response (e.g. “what was the identity of T2?”). In Experiment 2 we examined whether also non-conscious predictions might lead to subtle changes in subjective visibility reports. It was found earlier that predictions cannot be implemented by non-conscious stimuli (Meijs et al. 2018). That is, the benefit of prediction on T2 detection was dependent on correct T1 identification. Yet, as in this study, T2 was presented 275 ms or 917 ms after T1, at lag 3 and 10, respectively, it is possible that non-conscious predictions no longer exerted any effect by the time T2 was presented, given the weaker and more fleeting nature of non-conscious stimulus processing (Dehaene et al., 2006; Greenwald et al., 1996; van Gaal & Lamme, 2012; Van Vugt et al., 2018). Therefore, we used the same AB task as Meijs et al., but critically, T2 could now follow T1 much quicker as well, at lags 1, 2, and 10. Importantly, T1 prediction effects were measured both after correct T1 identification (“T1 seen”) and T1 missed trials (“T1 unseen”), which thus allowed us to address the question whether consciousness is necessary for prediction signaling (at least for the type of predictions studied here).

### 3.1 Methods and materials

#### 3.1.1 Participants

Forty students recruited from the University of Amsterdam participated in Experiment 2. We used a slightly larger sample size than in Experiment 1 because the effect size for non-conscious predictions was unknown. However, we have verified post-hoc that based on the sample size of 40 participants, we had 80% chance of detecting a non-conscious effect of prediction that is as small as η^2^=0.05.

All participants were right-handed, reported normal or corrected-to-normal vision, and had no history of a psychiatric or neurological disorder. All participants gave written informed consent prior to the start of the experiment and received research credits or money (10 euros per hour) for their participation. The full sample tested in Experiment 2 (11 males, mean age=20.41, SD=2.75) was included in the final analyses. The study was approved by the ethical committee of the Department of Psychology of the University of Amsterdam.

#### 3.1.2 Materials

The same materials as in Experiment 1 were used in Experiment 2.

#### 3.1.3 Task and Design

The task design and stimuli used in Experiment 2 were identical to Experiment 1 except for the following changes. On T1-present trials, T2 appeared at either lag 1 (92 ms), lag 2 (184 ms) or lag 10 (971 ms) after T1, with each lag being equally likely in each prediction condition. Second, T1 duration was titrated for each subject so that they each correctly reported T1 on approximately 65% of trials. The initial duration of T1 (92 ms) was adjusted after each experimental block depending on T1 identification performance. T1 duration was decreased for 1 frame (8 ms) when T1 identification performance exceeded 75% correct and was increased for 1 frame when performance would drop below 55% correct identifications. To make sure that T1 would not deviate too much from the duration of other RSVP stimuli, we kept it within the range of 42-142 ms (+/- 50 ms different from the normal RSVP item duration of 92ms). T1 was shown in white without placeholders (see Fig 2A). At the end of each trial, participants reported which T1 letter they saw by typing in H or G, or nothing when they did not see a T1, and then gave a rating of their perceptual experience of T2 on the PAS (PAS; 1-No experience; 2-A vague experience; 3-An almost clear experience; 4-A clear experience). Response measurement was thus identical to Experiment 1, except that the order of T1 and T2 responses was reversed.

**Figure 2.**
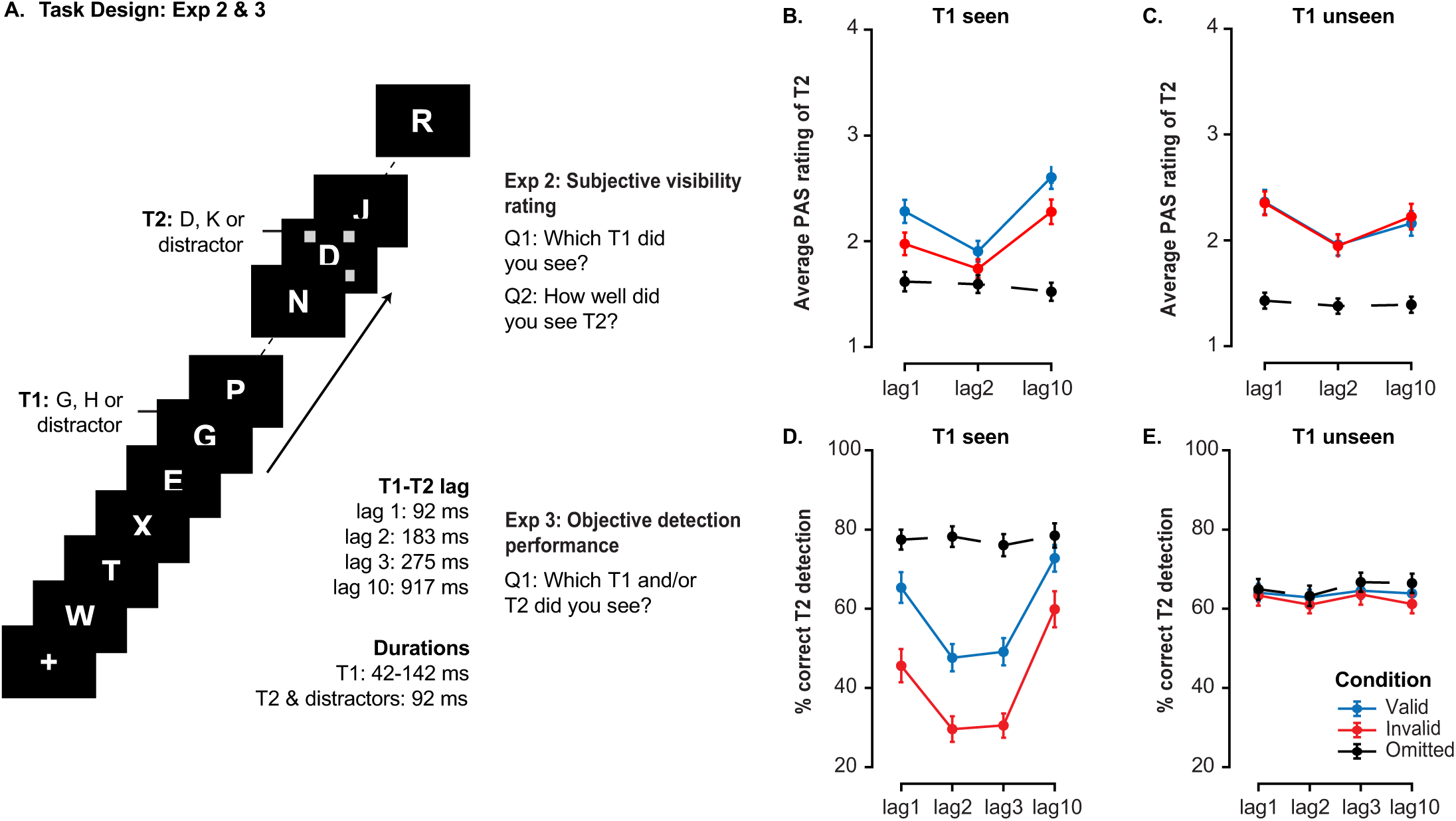
Task design and behavioral results of Experiment 2 and 3. **A)** The trial structure of the AB task used in Experiments 2 and 3. On each trial, we showed an RSVP stream of letters in which predefined target letters needed to be detected and reported at the end of the stream. The visibility of the first target (T1: G or H) was staircased so that the target was identified with approximately 65% accuracy. T1 always appeared at the fifth position in the stream. The second target letter (T2: D or K) could appear at one of three lags after T1 in Experiment 2 (lags 1, 2 and 10) and at one of four lags in Experiment 3 (lags 1, 2, 3 and 10). T2 was always surrounded with placeholders. On 20% of the trials, a random distractor letter was presented instead of T2 (T2 omitted trials). The identity of T1 predicted the identity of the T2 target letter with 75% validity, in the case a T2 was presented. This introduced validly and invalidly predicted T2s by seen and unseen T1s. At the end of each trial, participants needed to provide two answers. In Experiment 2, they needed to rate their subjective visibility of T2 using the PAS and then type in which T1 letter they had seen. In Experiment 3 participants were asked to type in which T1 and/or T2 they saw. **B, D**) PAS ratings (B) and percentage of correct T2 target discrimination (D) at each T2 lag for validly and invalidly predicted T2s and for T2 omitted trials on which T1 was correctly identified (T1 seen). When T1 was seen, validly predicted T2s were rated as more visible (panel B; Exp 2) and were detected more often (panel D; Exp 3) than invalidly predicted T2s. **C & E**) PAS ratings (C) and percentage of correct T2 target discrimination **(E)** at each T2 lag for validly and invalidly predicted T2s and for T2 omitted trials when T1 was missed (T1 unseen). Prediction did not modulate T2 visibility (panel C; Exp 2) or discrimination rate (panel E; Exp 3) at any of T2 lags when T1 was not seen. Error bars represent SEM.

#### 3.1.4 Procedure

Each participant completed two 2-hours-long sessions. Sessions were carried out on two different days. In session one, participants completed 3 practice blocks, followed by 15 experimental blocks, each block consisting of 60 trials in total. The second session consisted of 18 experimental blocks of 60 trials each. Everything else regarding the procedure of Experiment 2 was identical to Experiment 1.

### 3.2 Data Analysis

In Experiment 2, we tested whether average T2 PAS ratings differed significantly between validly and invalidly predicted T2s and critically, whether that depended on T1 visibility and/or T1-T2 lag. We computed average T2 PAS ratings using T1-present trials, separately for trials in which participants correctly identified T1 (T1-seen trials) and for trials in which T1 was missed (T1-unseen trials), separately for each lag, each prediction condition and for T2-omitted trials. Trials on which no response was given within the 4 seconds response timeout duration (on average, 3.29% of no T1 response trials and 6.1% of trials with no T2 ratings) or an impossible response was given (two T1 targets were reported on 0.03% of all trials) were excluded from the analyses.

Average T2 PAS ratings were entered into a 2 × 3 × 2 repeated-measures ANOVA with the within-subject factors T1 visibility (T1 seen, T1 unseen), lag (Lag 1, Lag 2, Lag 10) and prediction (valid, invalid). We also examined effects of T1 predictions on the PAS ratings separately for T1-seen and T1-unseen trials using two 3 × 2 repeated measures ANOVA with the within-subject factors lag (Lag 1, Lag 2, Lag 10) and prediction (valid, invalid). Significant main and/or interaction effects were followed-up by paired-sample *t*-tests. For all effects, we quantified the strength of evidence for the null hypothesis by computing a Bayesian equivalent of the same test (Wagenmakers et al., 2018).

### 3.3 Results

Our T1 duration staircasing procedure was successful in that average T1 accuracy was about 65% (mean=64.56%, SD=2.91%; T1 false alarm rate was 11.89%, suggesting that participants did not guess excessively). Trials in which T1 was correctly identified, were categorized as T1-seen trials, while T1-miss trials were considered as T1-unseen trials. T1 duration slightly differed between T1-seen (103.2 ms) and T1-unseen trials (101.4 ms, difference: t_39_=-5.3, p<.001, d=0.84, BF_01_=2.58×10^−4^), indicating that T1 visibility was associated with T1 duration. T1 duration however, did not influence whether T2 was seen or not (T2-present trials rated > 1 on the PAS scale, mean T1 duration = 101.85 ms; T2-present trials rated as 1 on the PAS scale, mean T1 duration=101.71 ms, t_39_=-0.145, p=.89, d=0.02, BF_01_=5.80).

Behavioral results are displayed in Figure 2. A repeated measures ANOVA on T2 PAS scores, including the factors T1 Visibility, Lag and Prediction validity revealed that T2 PAS ratings did not significantly differ between T1-seen and T1-unseen trials (F_1,38_=0.215, p=.65, η^2^=0.005, BF_01_=7.75), however not at all T2 lags (T1 Visibility × Lag: F_2,78_=30, p<.001, η^2^=0.44, BF_excl_=1.23×10^−6^). That is, PAS ratings were significantly higher for T1-unseen vs. T1-seen trials at lag 1 (t_39_=-2.64, p=.01, d=-0.42, BF_01_=0.28) and significantly higher for T1-seen vs. T1-unseen trials at lag 10 (t_39_ = 4, p<.001, d=0.63, BF_01_=0.01), but there was no difference at lag 2 (t_39_=-1.85, p=.07, d=-0.29, BF_01_=1.25). There was also a clear AB effect, indicated by a main effect of Lag (F_2,76_=32.19, p<.001, η^2^=0.45, BF_01_=1.81×10^−19^). Furthermore, validly predicted T2s were overall rated as more visible than invalidly predicted T2s (main effect Prediction; F_1,38_=17.28, p<.001, η^2^=0.31, BF_01_=0.13, no interaction with Lag: F_2,76_=0.63, p=.54, η^2^=0.02, BF_excl_=17.76). Crucially, the size of this prediction effect was modulated by T1 visibility (F_1,38_=35.37, p<.001, η^2^=0.476, BF_excl_=0.002), reflecting the fact that T1 prediction effects were only present in T1-seen but not T1-unseen trials (Figure 2B). The three-way interaction between all factors was also significant (F_2,76_=4.2, p=.019, η^2^=0.1, BF_01_=4.25), suggesting an additional influence of T1-T2 temporal interval, which we will unpack in more detail below based on the follow-up repeated measures ANOVAs we ran separately for T1-seen and T1-unseen trials.

Follow-up analyses revealed that predictions affected T2 visibility on T1-seen trials (F_1,39_=38.22, p <.001, η^2^=0.495, BF_01_=3.29×10^−4^), but not on T1-unseen trials (F_1,39_*=*0.36, p=.55, η^2^=0.009). A Bayesian equivalent of this analysis also yielded substantial support for the absence of a main effect of prediction on T1-unseen trials (BF_01_=6.61). The planned comparisons between T2 PAS ratings in valid and invalid conditions at each individual lag on T1-unseen trials also confirmed that non-conscious predictions effects were absent, even for the two shortest lags (lag 1: t_39_=0.03, p=.98, d=0.005, BF_01_=5.86; lag 2: t_39_=0.09, p=.93, d=0.014, BF_01_=5.84; lag 10: t_39_=-1.45, p=.16, d=-0.23, BF_01_=2.23). These results therefore confirm that non-conscious prediction effects, even when taking into account the quickly decaying nature of non-conscious processes, are not apparent at the shortest temporal lags. As such, they extend results from Meijs et al. (2018) who also showed that only consciously perceived T1s generated predictions biased conscious access, but only included longer-lag conditions (∼300 ms). Moreover, the size of the prediction effect on T2 PAS ratings in T1-seen trials was modulated by T2 lag (F_2,78_= 3.35, p=.04, η^2^=0.08, BF_excl_=4.33), although the effect was present for all lags (lag 1: t_39_ = 3.74, p<.001, d=0.59, BF_01_=0.02; lag 2: t_39_=4.61, p<.001, d=0.73, BF_01_=0.002; lag 10: t_39_ =6.84, p<.001, d=1.08, BF_01_=2.6×10^−6^).

Interestingly, in both T1-seen and T1-unseen trials, we observed lag-1 sparing and the AB (main effect of Lag on T1-seen trials: F_2,78_*=*47.95, p<.001, η^2^=0.55, BF_01_=1.26×10^−19^; main effect of Lag on T1-unseen trials: F_2,76_*=*17.29, p<.001, η^2^=0.31, BF_01_=4.45×10^−10^), suggesting that T1 awareness is not a necessary condition for these effects to occur. Specifically reflecting lag-1 sparing, T2s at lag 1 were rated significantly more visible than T2s presented at lag 2 (lag 1 vs. lag 2 seen: t_39_=8.75, p<.001, d=1.38, BF_01_=9.49×10^−9^; lag 1 vs. lag 2 unseen: t_39_=8.02, p<.001, d=1.27, BF_01_=7.95×10^−8^). Further, T2 ratings for lag 2 trials were significantly lower than T2 ratings at lag 10 (lag 2 vs. lag 10 seen: t_39_=-8.77, p<.001, d=- 1.39, BF_01_=9.09×10^−9^; lag 2 vs. lag 10 unseen: t_39_=-3.28, p=.002, d=-0.52, BF_01_=0.07), revealing an AB, irrespective of T1 visibility (see also Meijs et al. 2018).

## 4. Experiment 3

The findings from Experiment 2 suggest that even when undetected, T1 can impact subsequent T2 processing at lag 1 and within the AB window, yet conscious access to T1 is a prerequisite for initiation of prediction-related facilitation of T2 visibility reports. In a final experiment, we test the possibility that non-conscious predictions can modulate conscious access, as indexed by objective T2 discrimination, rather than self-reported subjective visibility. Specifically, we determined whether non-conscious predictions, measured at lag 1 and lag 2 (cf. Experiment 2 of the current study) and at lag 3 and lag 10 (the two lags used in Meijs et al. (2018, 2019), facilitate conscious access which we operationalized as the T2 discrimination rate. We reasoned that T2 discrimination rate may be more sensitive to decision-related shifts that might occur as a consequence of prediction signaling, which our study at this point cannot exclude as one of the mechanisms that drive prediction effects (if subjects merely guess T2 identity, even when they have not seen it, they will likely guess the predicted T2: Meijs et al., 2018). Our aim in Experiment 3 was two-fold. First, we aimed to replicate original findings by Meijs et al. (2018) and secondly, we wanted to test the hypothesis that the temporal scale on which weaker non-conscious effects are measured might prove critical for observing non-conscious prediction effects on T2 accuracy.

### 4.1 Methods and materials

#### 4.1.1 Participants

Forty students (28 female, mean age=22.1, SD=3.38) from the University of Amsterdam participated in Experiment 3 of this study. The rationale for choosing this sample size was the same as in Experiment 2. All participants were right-handed, and reported normal or corrected-to-normal vision and no history of a psychiatric or neurological disorder. All participants gave written informed consent prior to the start of the experiment and received research credits or money (10 euros per hour) as compensation. The study was approved by the ethical committee of the Department of Psychology of the University of Amsterdam.

#### 4.1.2 Materials

We used the same materials as in the previous two experiments.

#### 4.1.3 Task and Design

In two sessions, participants performed the AB task in which on each trial they needed to identify two target letters (T1 and T2) embedded in a stream of distractor letters. The task design of the current experiment was identical to Experiment 2 except for two changes. First, we added lag 3 (275ms) to the task design so that T2s could be presented at lag 1, 2, 3 and 10. Second, instead of using the PAS scale as our dependent measure, as in the original Experiment 3 by Meijs et al. (2018), we used T2 discrimination rate. Participants were asked to make a forced-choice judgement about the T2 letter identity by typing in this letter, after having made a forced-choice judgement about the T1 letter identity.

#### 4.1.4 Procedure

Each participant finished two 1.5h sessions on two different days. In both sessions, participants received detailed written and verbal instructions. All participants were explicitly informed about the predictive relationship between T1 and T2 and were told to use this information throughout the task. In the first session, participants first completed 3 practice blocks to get familiar with the task, and then another 15 experimental blocks. All blocks consisted of 60 trials each. In the second session, participants completed 21 blocks of 60 trials each without the practice. After every three blocks, there was a longer forced break during which the experimenter went into the testing cubicle to assure that participants took a break. Participants could also take shorter, self-timed breaks in between other blocks. At the end of each block, participants received written feedback about their performance (percentage of T1 and T2 misses and false alarms).

### 4.2 Data Analysis

T2 discrimination rate was computed separately per lag and prediction condition as the percentage of trials in which participants correctly reported the T2 letter identity when T2 was presented. On valid/invalid trials, T2 discrimination rate was computed on trials on which T1 was also correct. On omissions, trials were classified as correct when participants did not type in a T2 when T2 was not presented on T1-correct trials. Trials on which no T1 response was given within the 4 second response timeout duration (3.26% trials) or on which an impossible response was given, such as when two T1 or two T2 letters were typed in (0.6% and 0.22% trials, respectively), were excluded from the analyses.

In an attempt to minimize potential contributions of response bias to our results, we additionally performed a control analysis in which we used a measure of T2 detection performance (seen/miss) irrespective of T2 correctness (D/K) (Meijs et al., 2018). For that reason, this measure is independent of participants’ biases towards reporting the more likely T2 letter after having seen a specific T1 letter, since specific letter expectation should not influence participants’ ability to determine whether a target was simply presented or not. To that end, using T1-correct and T2-present trials, we computed this control dependent measure by aggregating all responses indicating the presence of T2, regardless of whether the reported letter was correct or not, as T2 seen trials, and no-T2 responses as T2 miss trials, and then computed the proportion of seen T2s in valid and invalid conditions. To illustrate, in the valid condition, the letter G is followed by the letter D, while in the invalid condition the letter G is followed by the letter K. If participants are simply *biased* to report the letter D based on the letter G even when they in fact did not see it, then there should be no difference between valid and invalid prediction conditions, since both conditions would contain roughly the same proportion of T2-seen responses.

T2 percentage correct discrimination was submitted to a 2 × 4 × 2 repeated-measures ANOVA with the within-subject factors T1 visibility (T1 seen, T1 unseen), Lag (Lag 1, Lag 2, Lag 3, Lag 10), and Prediction (valid, invalid). To follow up on this main analysis, we separated the analyses depending on T1 visibility. Using a 4 × 2 repeated measures ANOVA with the within-subject factors Lag (Lag 1, Lag 2, Lag 3, Lag 10) and Prediction (valid, invalid), separately for T1-seen and T1-unseen trials, we tested whether conscious and/or non-conscious predictions modulate conscious access to T2 presented at different lags. All effects were followed up by a Bayesian equivalent of the same test in order to evaluate the strength of evidence for the H_0_ (Wagenmakers et al., 2018).

### 4.3 Results

T1 identification rate was 65.1% (SD=3.72%; T1 false alarm rate was 14.93%, which suggests that participants did not guess excessively) and T1 duration did not differ between T1-seen vs. T1-unseen trials (although there was a trend, T1-seen=103.05 ms, T1-unseen=102.36 ms, t_39_=1.99, p=.054; d=0.31, BF_01_=0.99). A repeated measures ANOVA including the same 3 factors as in Experiment 2, but with T2 discrimination rate rather than T2 PAS rating as the dependent measure revealed largely similar effects. A robust AB was observed, as reflected by a main effect of Lag (F_3,117_=62.34, p<.001, η^2^=0.62, BF_01_=2.17×10^−9^), irrespective of prediction (Lag × Prediction: F_3,117_=0.85, p=.471, η^2^=0.02, BF_excl_=58.04). Replicating Meijs et al. (2018), predictions did increase T2 discrimination rate overall, with validly predicted T2s being identified more often than invalidly predicted T2s (F_1,39_=52.61, p <.001, η^2^=.574, BF_01_=2.98×10^−7^), and the size of this effect depended on T1 visibility (Prediction × T1 Visibility: F_1,39_=38.16, p<.001, η^2^=.495, BF_excl_=1.25×10^−7^). T2 discrimination was overall significantly lower after T1 was seen compared to when T1 was missed (F_1,39_=11.49, p = .002, η^2^=.228, BF_01_=1.64×10^−14^), especially at short lags (T1 visibility × Lag: F_3,117_=55.2, p<.001, η^2^=0.59, BF_excl_=1.6×10^−14^). Finally, as in Experiment 2, predictions influenced T2 discrimination differently across lags depending on T1 visibility (T1 Visibility × Lag × Prediction: F_3,117_=3.124, p=.029, η^2^=0.07, BF_excl_=17.34). In order to investigate this interaction effect in more detail, we followed this analysis up with two separate repeated measures ANOVAs, each testing the prediction effect on T2 discrimination rate at 4 different lags, separately for T1-seen and T1-unseen trials.

On T1-seen trials, T2 was detected more frequently when it was validly versus invalidly predicted (F_1,39_=48.94, p<.001, η^2^=.557, BF_01_=5.88×10^−14^). The size of this effect was modulated by lag (F_3,117_=3.445, p<.019, η^2^=0.08, BF_excl_=8.39), being numerically largest at lag 3 (t_39_=6.82, p<.001, d=1.08, BF_01_=2.73×10^−6^), although the effect of prediction was significant at each lag (all p’s<.001, all d’s>0.87, all BF_01_<3.97×10^−5^). On T1-unseen trials, prediction also seemed to affect T2 discrimination rate (F_1,39_=4.68, p=.037, η^2^=0.11), irrespective of lag (Lag × Prediction: F_3,117_=0.29, p=.833, η^2^=0.01, BF_excl_=22.16). However, a Bayesian analysis revealed stronger evidence for the null hypothesis, although anecdotal (BF_01_=1.74, in favor of the absence of a prediction effect) and planned pair-wise comparisons between valid and invalid conditions revealed that prediction did not affect T2 identification at any of four lags (all p’s>.136, all d’s<0.24, all BF_01_>2.03). Further, our control analyses carried out using the percentage of T2-seen responses as the dependent measure (i.e., disregarding T2 identity) confirmed that predictions affected T2 detection (irrespective of the correctness of this response) when T1s were seen (F_1,39_=44.47, p<.001, η^2^=.533, BF_01_=7.9×10^36^), however, not when T1 was unseen (F_1,39_=2.06, p=.16, η^2^=.05). Again, a Bayesian analysis revealed that evidence in favor of the null hypothesis was stronger than evidence in favor of a prediction effect on T1-unseen trials, although again evidence was anecdotal (BF_01_=2.72). Overall, these analyses suggest that a main effect of prediction after an unseen T1 is unlikely, and if anything, if present it may mainly reflect a higher guess rate on valid trials compared to invalid trials when T2 identification is the dependent measure of performance.

Overall, T2 discrimination differed across lags, as revealed by the main effect of lag. T2 performance on T1-seen trials showed the classical AB and lag-1 sparing. On T1-seen trials, there was a significant drop in T2 discrimination performance on lags 2 and 3 in comparison to the late lag 10 (main effect of Lag: F_3,117_= 81.75, p<.001, η^2^=0.68, BF_01_=1.02×10^−25^, lag 2 vs. lag 10: t_39_=-11.37, p<.001, d=-1.8, BF_01_=8.11×10^−12^; lag 3 vs. lag 10: t_39_=-10.8, p <.001, d=-1.71, BF_01_=3.44×10^−11^). Moreover, T2 discrimination performance was spared at lag 1 in contrast to other two short lags 2 and 3 (lag 1 vs. lag 2: t_39_=11.45, p<.001, d=1.81, BF_01_=6.63×10^−12^; lag 1 vs. lag 3: t_39_=7.55, p<.001, d=1.19, BF_01_=3.1×10^−7^). On T1 unseen trials the main effect of Lag was not significant (F_1,39_=1.128, p=.341, η^2^=0.028, BF_01_=12.78).

In conclusion, our results replicate and extend findings by Meijs et al. (2018), who showed that predictions increase the likelihood of conscious access, but only when predictions are consciously implemented. Using a wider temporal range between T1 and T2, that included shorter time intervals that could accommodate short-lived non-conscious predictions, we found no indication that predictions which are not consciously accessed facilitate conscious access.

## 5. Discussion

The current study addressed two outstanding questions concerning the relationship between prediction and consciousness. First, we examined if predictions can bias perception and increase subjective visibility reports of stimuli. Second, we determined if non-conscious predictions may selectively affect perception at short time scales, assuming the quickly-decaying nature of non-conscious processes (Dehaene et al., 2006; Greenwald et al., 1996; Van Vugt et al., 2018). To this end, in three experiments, we used an AB task in which the identity of T1 predicted which T2 target was most likely to appear (cf. Meijs et al., 2018, 2019). We determined whether predictions induced by seen and unseen T1s affected subjective measures of awareness (Exp 1 & 2) and/or objective discrimination performance of T2s (Exp 3). Experiment 1 showed that T2’s that confirmed predictions were reported to be perceived more clearly than T2s that violated predictions. In Experiment 2, we replicated and extended this finding by showing that prediction effects on subjective perceptual report depended on conscious access to the prediction-generating stimulus (i.e., T1), even at short time scales. In Experiment 3, we replicated the results of Experiment 2 but with objective T2 discrimination performance as the dependent variable. Based on these results, we conclude that predictions modulate subjective perceptual report of visual stimuli and the ability to consciously access them, but that this may critically depend on whether the prediction-initiating stimulus itself has been consciously accessed or not. Note that correct report was used as a measure of conscious access, but that, nevertheless, on some proportion of T1-seen trials, there might have been some T1 correct *guesses*, although T1 guess rate was found to be rather modest in all experiments. When signaled by unseen stimuli, (non-conscious) predictions did not systematically alter subsequent subjective visibility reports or discrimination performance, even though we measured their potential effects on a sufficiently short temporal scale to account for their alleged weaker and more fleeting nature. Further, participants always performed the AB task in two separate experimental sessions on different days, allowing for T1-T2 predictive associations to be formed. The fact that we did not consistently observe prediction effects on conscious access when T1 was unseen, even after such practice and training, extend previous findings (Bekinschtein et al., 2009; Meijs et al., 2019; Meijs et al., 2018; Strauss et al., 2015) that demonstrate a tight-knit relationship between predictions and consciousness.

In the first two experiments, we used the PAS scale to measure prediction effects. This scale emphasizes subjective perceptual aspects of awareness (Overgaard et al., 2006; Ramsøy & Overgaard, 2004) that might escape classical dichotomous (e.g., seen/unseen) measures or discrimination responses. The PAS scale has been shown to be sensitive to subtle changes in subjective visibility, even when these do not amount to a quantifiable effect on response correctness. For example, Overgaard et al., (2004) applied TMS bilaterally over the mid temporal regions in the ventral stream while participants viewed simple figures of different colors shown at different spatial positions. The authors found that TMS did not impair correctness of participants’ responses concerning different stimuli features, even though the participants experienced a decrease in subjective visibility when TMS was delivered 110-120 ms after stimulus presentation. This finding suggests that the PAS scale may capture certain aspects of reported awareness that are not always expressed in performance measures (for similar reasoning see: Lau & Passingham, 2006). Still, being a measure that relies on report, the PAS scale conceivably does not circumvent the problem of internal shifts in the response criterion, which could, for instance, manifest as a tendency to report higher levels of visibility for validly versus invalidly predicted targets (i.e. lenient versus more stringent response criterion for valid versus invalid trials). Therefore, the PAS scale arguably does not measure perceptual effects alone. Nevertheless, the overall picture was consistent across our experiments: predictions modulated subjective visibility reports, but they can only be implemented by correctly reported stimuli. These findings corroborate previous findings that used objective performance measures (Meijs et al., 2018) and extend these findings by showing that non-conscious predictions also do not affect conscious access at shorter time scales.

Implementation of the brain’s predictive mechanism has been described at various spatial scales – at the level of neural populations (Bastos et al., 2012), laminar connections in human primary visual cortex (Kok et al., 2016), cortico-striatal (den Ouden et al., 2009; den Ouden et al., 2012; Van Schouwenburg et al., 2010) and cortico-cortical interactions (Gordon et al., 2017; Summerfield & De Lange, 2014; Summerfield & Egner, 2009; Wacongne et al., 2011). It has been proposed that at each scale, prediction implementation and prediction error signaling are enabled by feedforward-feedback interactions (Friston, 2009). At a cortical level, after an initial feedforward sweep of activation of sensory cortices, sensory neurons receive feedback from higher-level cortical regions, thus implementing cortico-cortico feedback loops at various levels of the visual processing hierarchy (e.g., Summerfield & De Lange, 2014). Interestingly, influential models of consciousness posit that exactly this mechanism, i.e., feedback loops spanning widely-distributed brain areas and enabling sustained activation and global availability of information, is also necessary for conscious access (Dehaene et al., 2006; Dehaene & Changeux, 2011).

Based on our results, it could be concluded that predictive processing based on arbitrary associations between stimuli requires high-level neural feedback, characteristic for full-blown conscious access (Boly et al., 2017, 2013; Dehaene & Changeux, 2011). Relatively local processing of reentrant nature confined to visual cortices, which characterizes pre-conscious processes (Dehaene et al., 2006), may not be sufficient for implementing such predictions, at least not when recently learned, as in the current study. Future studies will need to address whether this conclusion still holds when predictions become over-trained and whether in such cases predictions may recruit different brain areas. It is also still an open question whether different types of prediction priors, for instance contextual and environmental priors acquired through lifelong experience (Brodski et al., 2015; Chang et al., 2016), those accrued from higher-order statistical regularities (Summerfield et al., 2008; Wacongne et al., 2011) and priors based on arbitrary stimulus associations (e.g. Kok et al., 2013; Meijs et al., 2019, 2018) rely on distinct neural mechanisms and may be implemented at different levels of the processing hierarchy. It is possible that more hard-wired priors that are acquired through life-long learning do not require conscious access to exert effects on perception, while arbitrary associations between stimuli such as those in the present study, depend on consciousness as they need to be implemented at higher processing levels.

An interesting aspect of our findings was that predictions initiated by consciously accessed T1s modulated T2 visibility and discrimination performance both at short and long lags, thus irrespective of temporal dynamics of attention deployment characteristic for RSVP tasks (Nieuwenstein, Burg, & Potter, 2009; Olivers & Meeter, 2008). This would also therefore suggest that predictions did not interact differently with exogenous (e.g., quickly triggered by T1 within ∼100 ms) versus slower endogenous attentional deployments (∼300 ms after a cue) which are known to have different time-courses (Carrasco, 2011). Predictions impacted performance at lag 1 (sparing), during the AB and outside the AB (lag 10). Similar observations have been reported previously (Meijs et al., 2018). Assuming that attention is mostly attenuated during the AB and less so at longer lags, this suggests that prediction effects operate relatively independent of attentional deployment in this task (see e.g. Kok, Jehee, et al., 2012). Regarding the nature of the prediction effect observed in the current study, one might wonder about the direction of the prediction effect: whether valid predictions led to a benefit in T2 performance or whether invalid predictions impaired it. Based on the current task design we cannot arbitrate between these two possibilities, because we did not include a neutral condition (“no prediction”) representing a baseline to which T2 performance in T2 valid and invalid conditions could be compared. However, using a very similar paradigm to the one administered in the current study, Meijs et al. (2019) addressed this issue directly and found that T2 discrimination performance for valid T2 trials was higher than for neutral trials and performance on invalid T2 trials was lower than on neutral trials. This thus suggests that valid predictions facilitate performance and invalid predictions worsen performance. Another interesting aspect of our results is that we found attentional enhancement of T2s at lag-1 and deficits at lag-2 and lag3, even when T1 was not consciously accessed (Experiment 2). This suggest that attention might have nevertheless been captured by T1 even when it was not processed all the way up to the level of report. This finding is in line with two earlier studies showing the same attentional enhancement (Nieuwenstein et al., 2009) and attentional blink (Meijs et al. 2018) after a missed T1.

In conclusion, we find a facilitatory effect of conscious, but not non-conscious, predictions on subjective visibility reports when arbitrary predictions need to be implemented on a trial-by-trial basis. This may suggest that high-level feedback signals that characterize full-blown conscious access are required for implementing relatively novel associative predictions learned over short periods of time. We show that a lack of prediction effect for non-conscious stimuli under these circumstances cannot be explained by the general fleeting nature of non-conscious processes and therefore, in line with earlier work, these findings demonstrate a close relationship between these types of predictions and consciousness.

## Acknowledgments

We thank Natalie Dikken and Amy de Boer for their valuable help with data acquisition. We thank Erik L. Meijs for sharing the task and analyses code of his original study, which were adapted for Experiment 3 of the current study. We thank Timo Stein for his insightful comments on a previous version of this manuscript. This work was supported by grants from the Psychology Research Institute of the University of Amsterdam.

